# IL-6 and IL-27 negatively regulate CRTAM-expressing CD4^+^ T-cells associated with lymphoid-driven synovitis

**DOI:** 10.1101/2024.09.03.610972

**Authors:** Alicia Derrac Soria, Myles Lewis, Xiao Liu, Jason P Twohig, Federica Monaco, Sandra Dimonte, Ana Cardus Figueras, Aisling Morrin, Carol Guy, Benjamin C Cossins, Robert Benson, Robert Andrews, Barbara Szomolay, Ernest H Choy, Paul Garside, Muneerah Huwaikem, Marc P Stemmler, Simone Brabletz, Thomas Brabletz, Florian Siebzehnrubl, Neil P. Rodrigues, Brendan J. Jenkins, Costantino Pitzalis, Gareth W Jones, Simon A Jones

**Affiliations:** Division of Infection & Immunity, School of Medicine, Cardiff University, Cardiff, Wales, UK; Systems Immunity University Research Institute, Cardiff University, Cardiff, Wales, UK; Centre for Experimental Medicine & Rheumatology, William Harvey Research Institute, Queen Mary’s School of Medicine & Dentistry, London, UK; School of Cellular & Molecular Medicine, University of Bristol, Bristol, UK; Centre for Immunobiology, Institute of Infection, Immunity & Inflammation, University of Glasgow, Glasgow, Scotland, UK; School of Biosciences, The European Cancer Stem Cell Research Institute, Cardiff University, Cardiff, Wales, UK; Department of Experimental Medicine, Nikolaus-Fiebiger-Center for Molecular Medicine, FAU University Erlangen-Nürnberg, Erlangen, Germany; South Australian immunoGENomics Cancer Institute (SAiGENCI), The University of Adelaide, Adelaide, South Australia, Australia

**Keywords:** Cytokines, gp130, STAT transcription factors, ectopic lymphoid structures, arthritis

## Abstract

Joint pathology in rheumatoid arthritis is heterogeneous, with histology providing evidence of fibroblast-driven, myeloid-driven, and lymphoid-driven synovitis. However, the immuno-modulatory pathways underlying their development remain unclear. Profiling synovial tissues from rheumatoid arthritis patients and mice with antigen-induced arthritis, we identified a subset of synovial infiltrating CD4^+^ T-cells expressing CRTAM (class-I MHC-restricted T-cell-associated molecule). In human synovial biopsies, *CRTAM* correlated with the expression of effector cytokines (*IL21*, *IFNG*), chemokine receptors (*CXCR3*, *CXCR4*, *CCR5*), granzymes (*GZMA*, *GZMB*, *GZMK*), and regulatory factors (*TIGIT*, *EOMES*, *BATF*) linked with T-cell-mediated immunity. Studies of antigen-induced arthritis showed that CRTAM^+^CD4^+^ T-cells accumulate in the inflamed synovium following disease onset. CRTAM^+^CD4^+^ T-cells were particularly abundant in synovial tissue from *Il27ra*^-/-^ mice displaying ectopic lymphoid-like structures. CADM1 (cell adhesion molecule-1), the endogenous ligand for CRTAM, was also expressed in human synovitis and synovial tissues from wild-type, *Il6ra*^-/-^, and *Il27ra*^-/-^ mice with antigen-induced arthritis. Cells expressing human CADM1 included synovial fibroblasts and subsets of monocytic and CD19^+^ cells. Considering the *ex vivo* regulation of CRTAM, we identified that activation of naïve CD4^+^ T-cell increased CRTAM expression. This induction was blocked by IL-6 and IL-27, with further studies identifying a role for STAT3 in controlling the CRTAM transcriptional repressor, ZEB1. These results provide insights into the cytokine control of CRTAM on CD4^+^ T-cells and support the involvement of CRTAM^+^CD4^+^ T-cells in lymphoid-driven synovitis.

## INTRODUCTION–

Studies of cytokines in immune-mediated inflammatory diseases (IMIDs) have pioneered advances in biological and targeted medicines that inhibit their signaling properties^1–3^. However, patients often show varying efficacies to certain drug classes that reflect the complex and heterogeneous nature of the underlying pathology^2,3^. In rheumatoid arthritis (RA), human synovial biopsies display a range of cellular and molecular hallmarks of disease that inform the classification of fibroblast-rich, myeloid-rich, and lymphoid-rich synovitis^4–6^. While the mechanisms steering these pathologies are unknown, mouse models and human transcriptomic data point toward cytokines acting *via* the Janus-activated kinase-signal Transducer and Activator of Transcription (Jak-STAT) pathway^4,7–11^.

The Jak-STAT pathway senses and interprets cytokine cues essential for cellular survival and functional identity^12–16^. Treatments for IMIDs with blockers of Jak-STAT signaling (e.g., oral Jak inhibitors and IL-6 antagonists) target STAT1 and STAT3 transcription factors as part of their mode of action^12–15^. These transcription factors display a complex inter-relationship that impacts leukocyte recruitment and activation, synovial hyperplasia, and joint erosion^9,17–21^. Investigating STAT1 and STAT3 activities in wild-type, *Il6ra*^-/-^ and *Il27ra*^-/-^ mice with antigen-induced arthritis (AIA), we established that these mice develop joint pathologies resembling the heterogeneity of synovitis in humans^4,7–11^. Appreciating the limitations of these models to the study of RA in humans, these genetic strains provide a first-up opportunity to define the molecular basis of disease heterogeneity in synovitis. Combining transcriptomic datasets from murine CD4^+^ T-cells, synovial tissues from mice with inflammatory arthritis, and human synovial joint biopsies, we aimed to identify gene signatures of Jak-STAT signaling that shape the course of cytokine-driven pathology. Studies described herein identify several gene signatures associated with lymphoid-like synovitis, including an immunoglobulin-like cell surface protein of the Nectin-like family of adhesion molecules (termed; Class I-restricted T-cell associated molecule; CRTAM, CD355)^22^.

CRTAM is primarily associated with the effector properties of activated NK cells, NKT cells, and CD8^+^ T-cells^23–28^. However, CRTAM is also seen on a subset of CD4^+^ T-cells following TCR activation^29,30^. CRTAM signals *via* cell-adhesion molecule-1 (CADM1; also known as Necl2), an endogenous membrane-associated ligand expressed on stromal cells. This interaction controls the cytotoxic and effector properties of lymphoid cells. For example, CRTAM^+^CD4^+^ T-cells display heightened IFNγ, IL-17A, and IL-22 expression and increased cell migration to inflammatory foci^30–33^. These activities require contact adhesion of CRTAM^+^ T-cells with CADM1-expressing cells^28^. Draining lymph nodes or ectopic lymphoid-like structures may represent sites for these interactions, with studies identifying regulatory roles for CRTAM in adaptive mucosal immunity against infection, resistance to autoimmune diabetes, and thymic development^24,25,30,34–36^. Confirming the expression of CRTAM by activated naïve CD4^+^ T-cells^29,33^, we now show that IL-6 and IL-27 negatively regulate CRTAM induction following TCR stimulation, with analysis of human RA and mice with AIA identifying CRTAM^+^CD4^+^ T-cells as a feature of lymphocyte-driven synovitis. We, therefore, propose the CRTAM-CADM1 signaling system as a potential determinant of effector T-cell responses linked with synovial autoimmune reactions.

## MATERIALS AND METHODS

### Mice–

Inbred wild-type mice were purchased from Charles River UK. The *Il27ra*^-/-^ mouse strain (line B6N.129P2-*Il27ra*^tm1Mak^/J) was from Jackson Laboratory and generated by genetic disruption of the extracellular fibronectin type III domain of IL-27R^37^. Inbred *Il6ra*^-/-^ mice were generated by disrupting exons 4, 5, and 6, encoding the IL-6 binding domain^38^. Mice with altered gp130 signaling were generated by a phenylalanine substitution of tyrosine-757 in the gp130 cytoplasmic tail (*gp130*^Y757F:Y757F^)^39,40^. Compound *gp130*^Y757F:Y757F^*:Stat3*^+/-^ mice were generated as described^39,40^. The generation of *Zeb1*^-/-^ mice is previously described^41^.

### Antigen-induced arthritis (AIA)–

Animal procedures for AIA were performed under approved UK Home Office licenses (PPL30/2928 and PB3E4EE13)^7–9,19,42^. Male mice were immunized (s.c.) with 100 μl methylated BSA (1 mg/ml emulsified in Complete Freund’s Adjuvant; CFA) and 160 ng *Bordetella pertussis* toxin (i.p.) (all from Sigma-Aldrich). Mice were re-challenged with methylated BSA (mBSA) and CFA (s.c.) one week later. Arthritis was induced on day-21 by administration (i.a.) of mBSA (10μl; 10 mg/ml) into the right knee joint. Synovial tissues were harvested on day-3 and day-10 following disease onset to reflect early and established forms of the disease^8^. For flow cytometric analysis of synovial CD4^+^ T cells, the inflamed synovium was dissected, digested in collagenase type IV (37°C, 1h) and cells recovered through a 40μm cell strainer (Greiner Bio-one)^10^.

### Histological assessments–

Knee joints from mice with AIA were decalcified before formalin fixation and paraffin embedding. Parasagittal serial sections of 7μm thickness were prepared and stained in hematoxylin, fast green, and safranin-O. Pathology was scored by 3 assessors blinded to study design for synovial infiltrate (0-5), synovial exudate (0-3), synovial hyperplasia (0-3), and cartilage and joint erosion (0-3)^7–9^. An Arthritis Index was obtained from accumulative scores. For immunohistochemistry, sections were rehydrated and treated with 10mM sodium citrate buffer containing 0.05% (v:v) Tween-20 (95°C, 40 min). Non-specific antibody binding was reduced by incubating sections with 10% (v:v) goat or swine serum (according to the secondary antibody). Endogenous peroxidase activity was blocked with 3% (v:v) H2O2. Immunohistochemistry was performed with antibodies to CD3 (Clone A0452, Dako), CRTAM (Clone D3A7; Vector Laboratories), and ZEB1 (polyclonal; Santa Cruz Biotechnology). Following secondary antibody labeling (E0431; Dako), immuno-staining was detected using the Vectastain ABC kit and diaminobenzidine (Vector Laboratories). Sections were counterstained with hematoxylin and imaged on a Leica DM2000 LED before quantification with Leica QWin software^10^.

### T-cell isolation–

Splenic murine CD4^+^ T-cells were isolated from spleens homogenized through a 40μm cell strainer (Greiner Bio-one)^8,10,38^. Recovered cells were washed and red blood cells lysed before negative selection of CD4^+^ T-cells by magnetic-activated cell sorting (MACS; Milteny Biotech). Naïve CD4^+^ T-cells were enriched by flow cytometry cell sorting using fluorochrome-conjugated antibodies against CD4, CD25, CD44, and CD62L on a FACS Aria II system (BD Biosciences). Isolated naïve CD4^+^ T-cells were identified as CD4^+^CD25^-^CD44^lo^CD62L^hi^ cells and displayed a purity of >95%.

### T-cell stimulation–

Naïve CD4^+^ T-cells (10^5^ cells/well) were seeded in supplemented RPMI-1640 medium into Nunclon delta surface 96-well U-bottom plates (Thermo Fisher Scientific)^8,10,38^. For TCR activation, cells were co-stimulated with anti-CD3 (1 μg/mL anti-CD3; clone 145-2C11, R&D systems) and soluble anti-CD28 (5 μg/mL; clone 37.51, eBioscience)^8,10,38^. Cells were treated with IL-6 (20 ng/mL), IL-12 (10 ng/mL), or IL-27 (10 ng/mL) as indicated. Recombinant cytokines were purchased from R&D Systems. Cells were grown at 37°C with 5% CO_2_.

### Flow cytometry–

Samples were acquired on a BD FACSCanto II or Attune flow cytometer using FACSDiva software (BD Biosciences) and analyzed with FlowJo v10 (Tree Star Inc, US)^8,10,38^. Cell numbers were quantified using counting beads (Life Technologies). Antibodies for the following antigenic markers were used: CD3 (Clone 145-2C11; BioLegend), CD4 (Clone RM4-5), CD25 (Clone PC61.5), CD44 (Clone IM7), CD62L (Clone MEL-14), CD8a (Clone 53-6.7), γ8TCR (eBioGL3), NK1.1 (Clone PK136; all from eBioscience), and CD355/CRTAM (Clone 11-5; BioLegend).

### Quantitative PCR–

Gene expression was quantified by TaqMan Gene Expression Assay (Applied Biosystems) using a barcoded MicroAmp Optical 384-well reaction plate (Applied Biosystems)^8,10^. Taqman oligonucleotide primers for *Crtam* (Mm00490300_m1), *Zeb1* (Mm00495564_m1), *Cadm1* (Mm00457551_m1), and *Actb* (Mm01205647_g1) were purchased from Thermofisher.

### Chromatin immunoprecipitation (ChIP)-qPCR–

Genomic DNA was extracted from activated naïve CD4^+^ T-cells (10^7^ cells/ml) following a 6-hour stimulation with IL-6 (20 ng/ml), IL-27 (10 ng/ml) or vehicle control. Protein-DNA complexes were cross-linked with formaldehyde, and cell nuclei recovered before sonicating into chromatin fragments of 150-600bp. ChIP was performed with 5μg of anti-STAT1 (sc-592) or anti-STAT3 (sc-482) antibodies (Santa Cruz Biotechnology). Isotype-specific IgG antibodies were used as controls. Antigen-antibody complexes were isolated using Protein-A/G magnetic beads (Pierce) before DNA extraction for qPCR (QuantStudio 12K Flex system) with oligonucleotide primer sequences mapped to gene promoters^10^. Customized primers were generated for promoters of *Irf1* (Forward strand: CCTTCGCCGCTTAGCTCTAC; Reverse strand: CCCACTCGGCCTCATCATT), *Socs3* (Forward strand: CTCCGCGCACAGCCTTT; Reverse strand: CCGGCCGGTCTTCTTGT), and *Zeb1* (Forward strand: CAAAGTGCGCAACTCGTC; Reverse strand: TCCTAGGCCAGGTTGAGC). The enrichment of promoter sequences was normalized to values obtained with isotype control antibodies and input controls and expressed as 2^^ΔCT^.

### Human samples and other repository datasets–

Details of the clinical cohorts and human synovial biopsy studies are provided here^4–6,43^. Transcriptomic data from the PEAC and R4RA studies and the *Accelerated Medicines Partnership* RA Arthroplasty phase-1 study are available (accession numbers E-MTAB-6141 and SDY998)^4,6,43^. Transcriptomic datasets from murine CD4^+^ T-cells are accessible here (E-MTAB-7682)^10^.

### Statistics–

Data analysis was conducted with GraphPad Prism 7 software (GraphPad Software Inc., La Jolla, USA). Student’s t-test determined the significance of two experimental groups with normal Gaussian distribution. For data that were not normally distributed a Mann-Whitney U-test was used. Statistical significance between groups of normally distributed data was tested using one-way ANOVA with Turkey’s multiple comparisons. Where experimental groups were compared across time points, a two-way ANOVA was used with a Bonferroni post-test. *P*<0.05 was considered statistically significant.

## RESULTS–

### Synovial CRTAM expression is associated with lymphoid-rich synovitis–

RNA-seq datasets from human synovial biopsies showed a differential expression of *CRTAM* in tissues characterized by histology as fibroblast-rich, myeloid-rich, and lymphoid-rich synovitis (**Fig-1A**)^4–6^. *CRTAM* was highly expressed in lymphoid-rich synovitis and showed positive correlations with disease activity scores and phenotypic markers of lymphocyte activation (**Fig-1B** & **C**). These included regulatory molecules (e.g., *CD2*, *CTLA4*, *SLAMF7*, *TIGIT*), chemokine receptors (e.g., *CXCR3*, *CXCR4*, *CXCR6*, *CCR5*), granzymes (e.g., *GZMA*, *GZMB*, *GZMK*), and transcription factors (e.g., *BATF*, *EOMES*) (**Fig-1C**). To verify these findings, we screened single-cell RNA-seq data of synovial biopsies reported by the *Accelerating Medicines Partnership*^43^. Lymphocytes defined by granzyme expression or the markers CCR7 or PD-1 showed high CRTAM expression (**Fig-1D**). Consistent with the enhanced expression of effector cytokines by CRTAM^+^ lymphocytes^30,33^, we identified positive correlations between synovial *CRTAM* and *IFNG,* and *CRTAM* and *IL21* in two patient cohorts (**Fig-1E**). Comparisons with *IFNA*, *IFNB*, *IL17A*, and *IL22* showed weaker correlations (**Fig-1E**). These data identify a link between synovial *CRTAM* and the effector properties of lymphoid cells in synovitis^8,36,44^.

**Figure-1.**
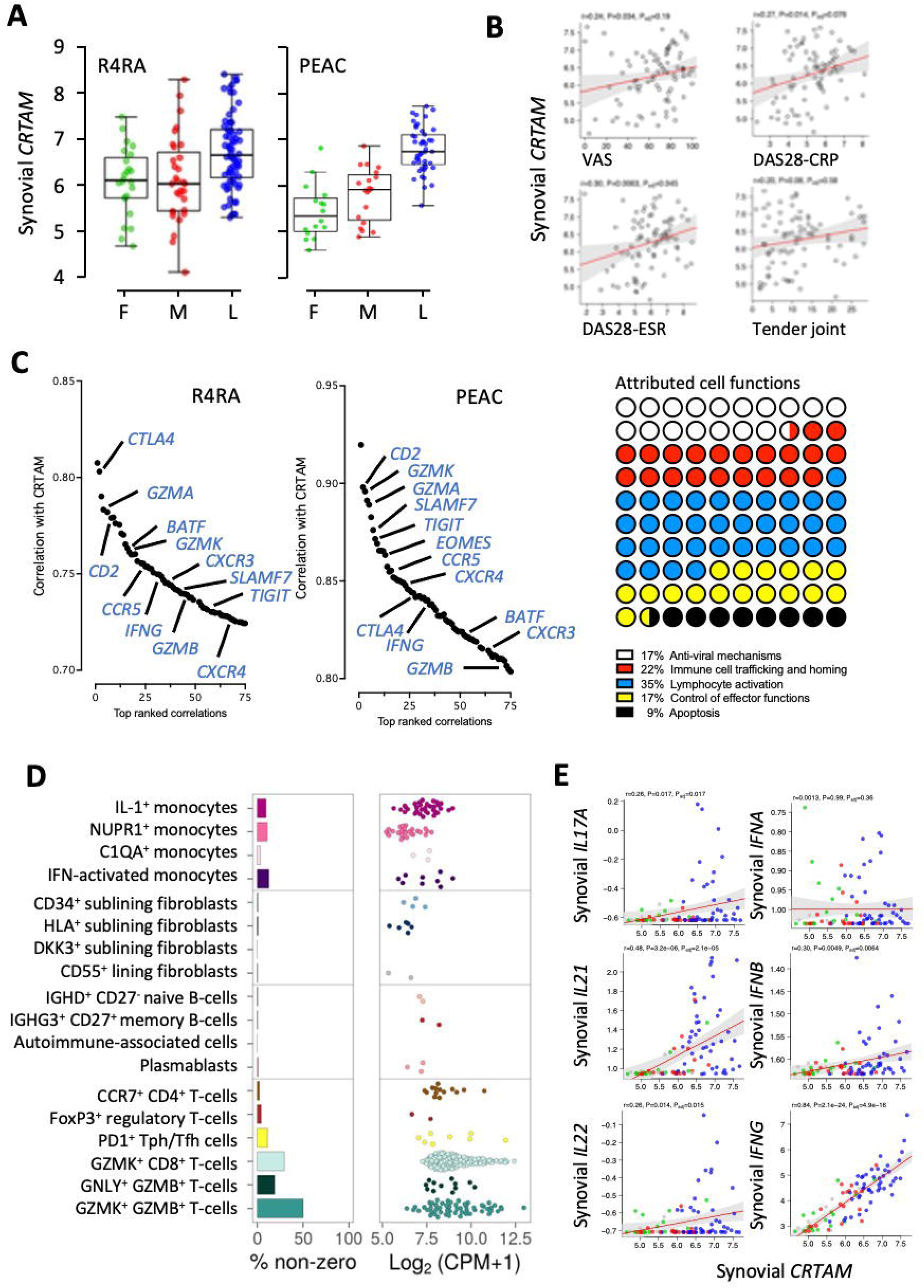
CRTAM expression is associated with lymphoid-rich synovitis. **(A)** *CRTAM* expression in synovial tissue acquired through a minimally invasive ultra-sound synovial biopsy from RA patients enrolled in the R4RA and PEAC studies. Synovial expression derived by bulk RNA-seq is shown for patients stratified according to synovial pathology (lymphoid, blue; myeloid, red; fibroid, green). **(B)** Analysis of RNA-seq data from synovial biopsies collected through the R4RA trial. Pearson correlations of synovial *CRTAM* against parameters of disease activity (DAS28 scores, tender joint scores, VAS). **(C)** Additional correlations with other inflammatory markers, ranking the Top 75 gene correlations from R4RA and PEAC. A breakdown of the biological functions associated with these genes is shown. **(D)** Classification of synovial *CRTAM* in scRNA-seq from the *Accelerating Medicines Partnership*^43^. **(E)** Pearson correlations of synovial *CRTAM* with the inflammatory cytokines *IFNG*, *IFNA*, *IFNB*, *IL17A*, *IL21*, and *IL22* (data from the R4RA study).

### Detection of synovial CRTAM*^+^* lymphocytes in mice with inflammatory arthritis–

We next evaluated the synovial expression of CRTAM in wild-type (*wt*), *Il6ra*^-/-^, and *Il27ra*^-/-^ mice with AIA (**Fig-2**). These mouse strains develop distinct forms of synovitis in response to AIA, with pathologies reflecting the heterogeneity of synovitis in RA^4,7–11^. In this regard, *Il27ra*^-/-^ mice with AIA developed a more severe form of pathology than *wt* mice, with *Il6ra*^-/-^ mice showing signs of early synovial fibroblast activation in the absence of an immune cell infiltrate (**Fig-2A**). Harvesting joint sections and synovial mRNA from non-challenged mice and mice with AIA, we monitored CRTAM expression following disease onset (on days 3 and 10 of AIA). Immunohistochemistry showed that CRTAM primarily co-localized with infiltrating CD3^+^ lymphocytes (**Fig-2B** & **2C**). CRTAM expression was also observed in the synovial lining (best illustrated by CRTAM staining in sections from *Il6ra*^-/-^mice with AIA). Analysis of synovial mRNA by qPCR confirmed an early induction of *Crtam* following AIA onset (i.e., day 3 of disease) (**Fig-2D**). Synovial *Crtam* was most pronounced in tissues from *Il27ra*^-/-^ mice, whereas *Il6ra*^-/-^ mice with AIA showed the smallest increase in *Crtam* (**Fig-2D**).

To identify CRTAM-expressing cells, synovial infiltrating leukocytes were extracted from the inflamed synovium of mice with AIA (at days 3 and 10) and analyzed by flow cytometry. Since *Il6ra*^-/-^mice lack a synovial infiltration in response to AIA, we limited our analysis to wild-type and *Il27ra*^-/-^mice. We observed increases in synovial CRTAM^+^CD3^+^ lymphocytes following arthritis induction (**Fig-2E**). These cells were predominantly CD4^+^ T cells (totaling 70-80% of the lymphocyte infiltrate) and were particularly evident in *Il27ra^-/-^* mice with AIA (**Fig-2E**).

**Figure-2.**
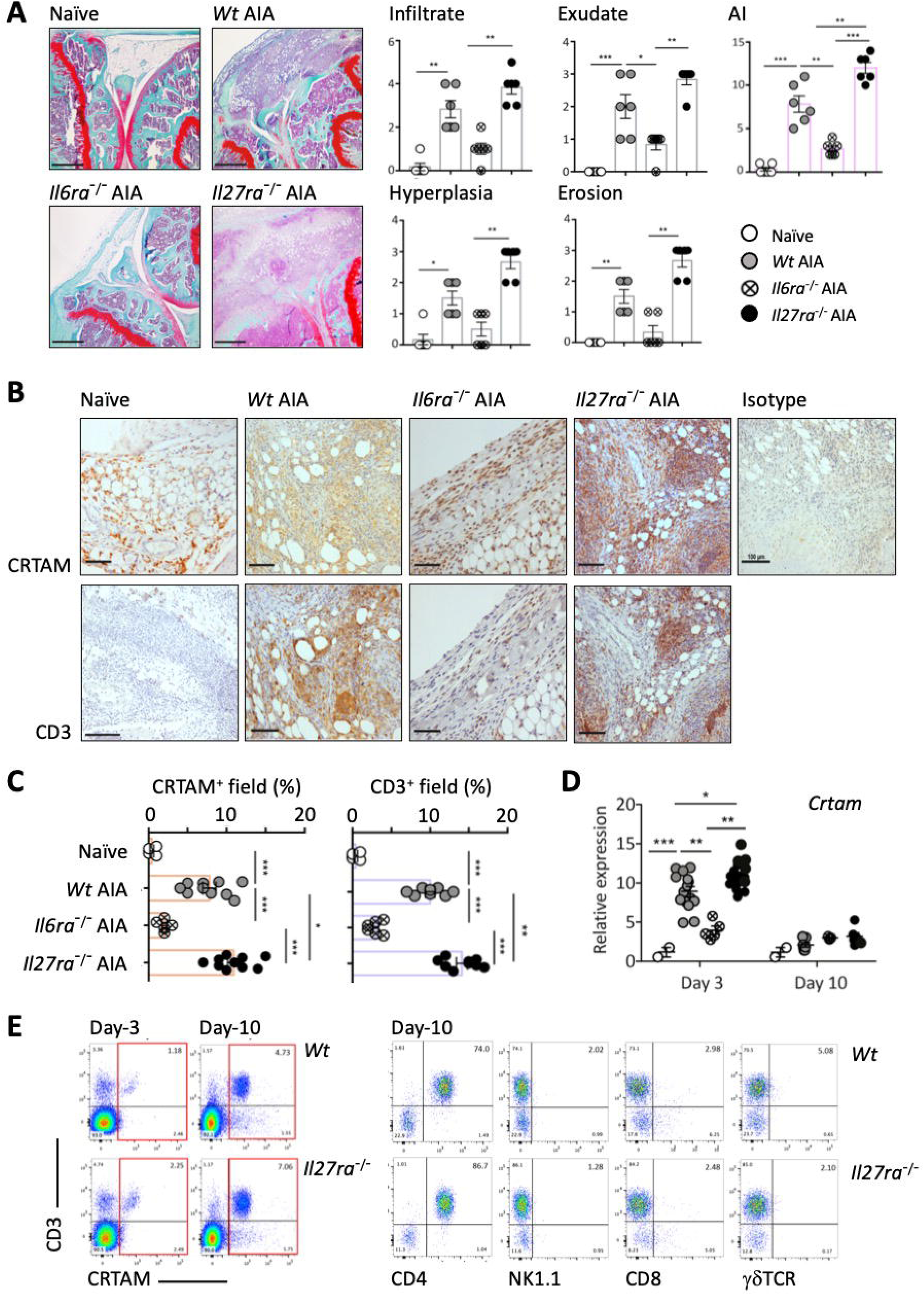
Identification of synovial CRTAM*^+^*CD4*^+^* T-cells in antigen-induced arthritis. **(A)** Antigen-induced arthritis (AIA) was established in wt, *Il6ra*^-/-^ and *Il27ra*^-/-^ mice (scale bar, 500μm). Histology scores for disease parameters are shown with an aggregate Arthritic Index (day-10 of AIA). Values show mean ± SEM (n=5-6 mice/group; \**P*<0.05; \*\**P*<0.01; \*\*\**P*<0.001). **(B)** Representative immunohistochemistry of knee joints taken on day-10 following arthritis onset (scale bar, 100μm). Staining is shown for CRTAM and CD3. **(C)** Association of CRTAM staining against that of CD3 in synovial tissues (mean ± SEM; n=5-11 mice/group; \**P*<0.05; \*\**P*<0.01; \*\*\**P*<0.001). **(D)** Quantitative PCR of synovial *Crtam* in mice with AIA (day-3 and day-10 of disease) and correlation of expression with histological scores of disease pathology (mean ± SEM; n=2-20 mice/group collected from three independent experiments; \**P*<0.05; \*\**P*<0.01; \*\*\**P*<0.001). **(E)** Flow cytometric analysis of CRTAM-positive cells in infiltrates extracted (day-3 and day-10 of disease) from the inflamed synovium of *wt* and *Il27ra*^-/-^ mice with AIA.

### Synovial expression of the CRTAM ligand CADM1–

Synovial *CADM1* remained comparable across all three forms of human synovitis and showed no correlation with parameters of disease activity (e.g., DAS28 scores, ESR, tender joint scores, and pain) (**Fig-3A** & **B**). We saw a similar profile of *Cadm1* expression following arthritis induction in mice, with synovial mRNA levels remaining comparable between *wt*, *Il6ra*^-/-^, and *Il27ra*^-/-^ mice (**Fig-3C**). These results suggest that CADM1 is less prone to inflammatory regulation and unaltered by synovial infiltrating leukocytes. Investigating single-cell RNA-seq data from the *Accelerating Medicines Partnership*, we identified sub-lining fibroblasts and IL-1β^+^ and NUPR1^+^ monocytes as synovial cells expressing CADM1 (**Fig-3D**). Thus, CRTAM and CADM1 may facilitate cellular interactions between stromal cells within the inflamed joint and synovial infiltrating lymphocytes.

**Figure-3.**
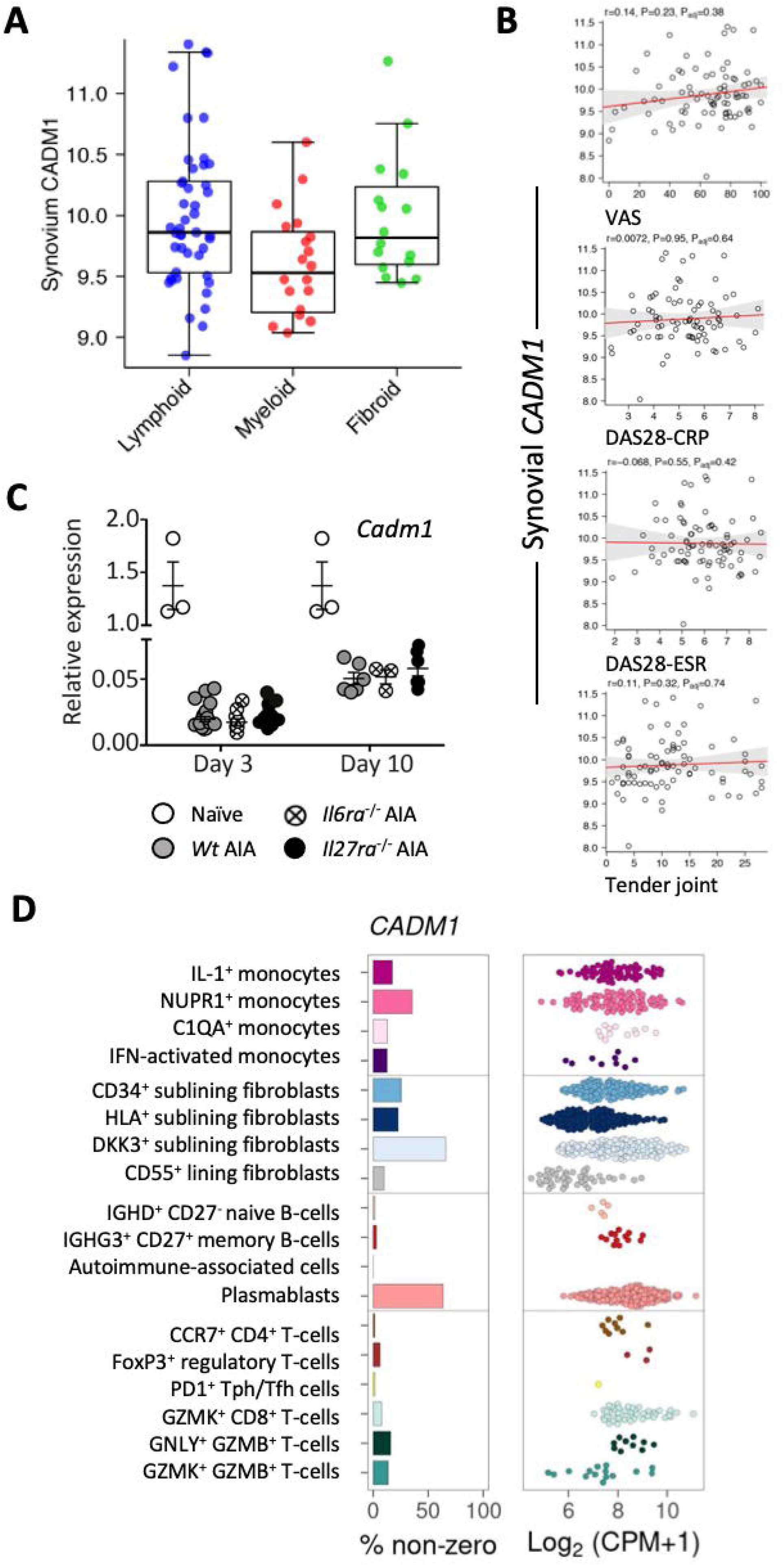
CADM1 is regulated in response to synovitis. **(A)** *CADM1* expression in human joint biopsies stratified according to synovial pathology (lymphoid, blue; myeloid, red; fibroid, green). Values indicate mean ± SEM (n=5-20 mice/group collected from three independent experiments; \**P*<0.05; \*\**P*<0.01; \*\*\**P*<0.001). **(B)** Pearson correlations of synovial CADM1 with DAS28 scores, tender joint scores, and VAS. **(C)** Quantitative PCR of synovial *Cadm1* in synovial tissue from wt, *Il6ra*^-/-^ and *Il27ra*^-/-^ mice with AIA (day-3 and day-10 of disease). **(D)** Synovial *CADM1* in scRNA-seq from the *Accelerating Medicines Partnership* datasets^43^.

### Cytokine control of CRTAM*^+^*CD4*^+^* T-cells–

To understand the regulation of CRTAM on CD4^+^ T-cells, we used multi-parameter flow cytometry to investigate temporal changes in CRTAM on activated naïve CD4^+^CD25^-^CD44^lo^CD62L^hi^ T-cells (**Fig-4**). Following co-stimulation with anti-CD3 and anti-CD28 antibodies, we identified a subset of activated naïve CD4^+^ T-cells expressing CRTAM (representing 15% of cells). Maximal expression was observed 18 hours after stimulation (**Fig-4A**). Reflecting on the increased incidence of synovial CRTAM^+^ lymphocytes in *Il27ra^-/-^* mice with AIA (**Fig-2**), we tested whether 20 ng/ml IL-27 affects CRTAM expression on activated naïve CD4^+^ T-cells. Tracking the longitudinal expression of CRTAM following anti-CD3 and anti-CD28 antibody stimulation, IL-27 reduced CRTAM expression by 50-60% (**Fig-4B**). No change in CRTAM expression was observed in response to IL-12 (**Supplementary Figure-1A**). Extending these findings to other gp130-activating cytokines, we assessed the activity of IL-6. Transcriptomic data from IL-6 stimulated naïve CD4^+^ T-cells (E-MTAB-7682) showed that IL-6 inhibited the expression of various genes induced by TCR activation (e.g., *Xcl1*, *Stap1*, *Lta*, *Art2b*, *Tnfsf14* and *Xdh*), including *Crtam* (**Supplementary Figure-1B**). The *ex vivo* activation of naive CD4^+^ T-cells endorsed these findings, with 20ng/ml IL-6 suppressing the proportion of CRTAM^+^CD4^+^ T-cells detected by flow cytometry (**Fig-4B**). The specificity of the *Crtam* inhibition by IL-6 and IL-27 was confirmed using splenic naïve CD4^+^ T-cells from *Il6ra*^-/-^ and *Il27ra*^-/-^ mice (**Fig-4C**). Thus, lymphokines regulate *Crtam* induction in response to naïve CD4^+^ T cell activation.

**Figure-4.**
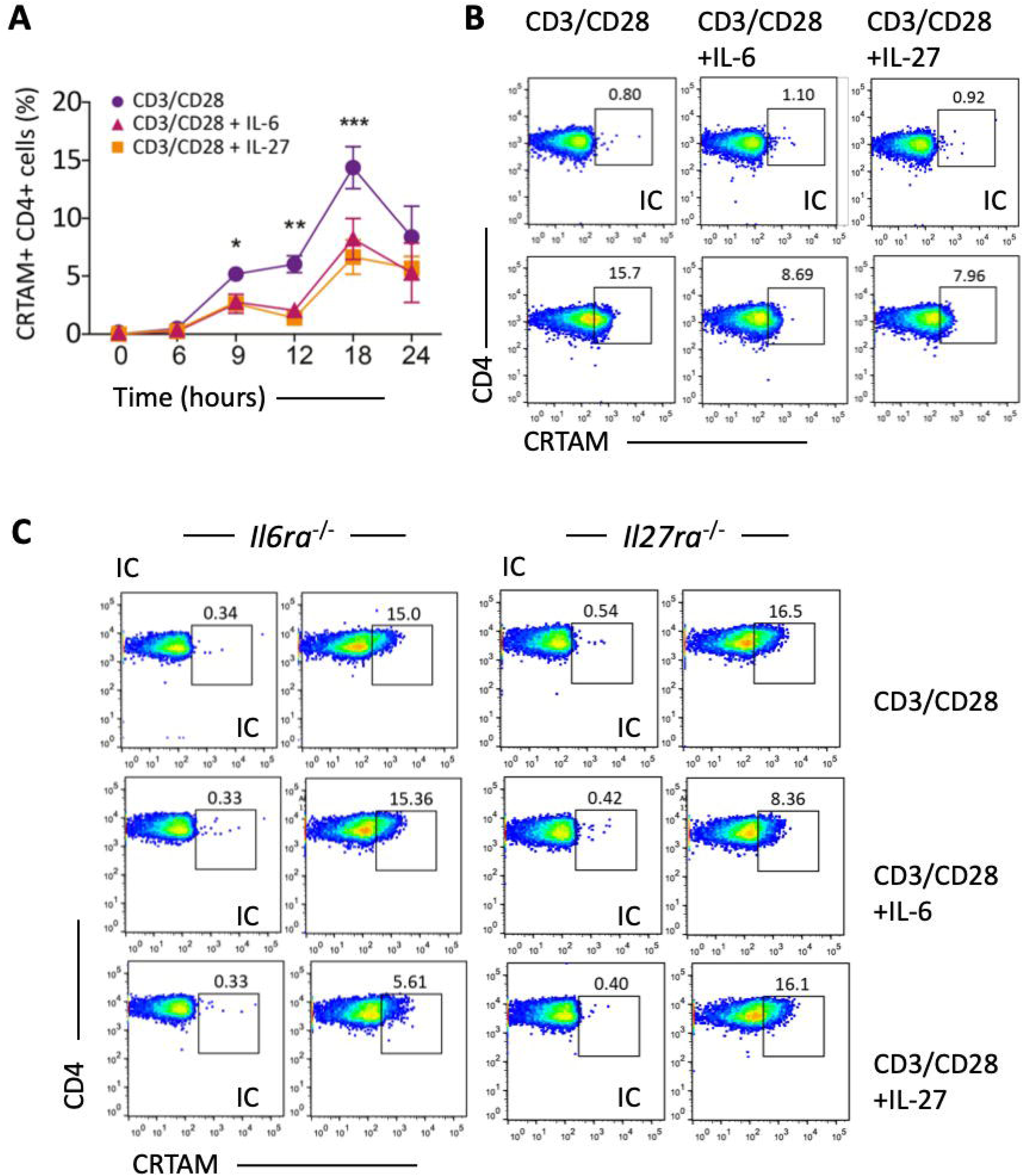
IL-6 and IL-27 blocks CRTAM expression on activated naïve CD4+ T-cells. **(A)** Splenic naïve CD4^+^ T-cells from *wt* mice were cultured with co-stimulatory anti-CD3 and anti-CD28 antibodies. Temporal changes in CRTAM^+^ CD4^+^ T-cells are shown following treatment with 20 ng/ml IL-6 or 10 ng/ml IL-27 (mean ± SEM from 3 experiments; \**P*<0.05; \*\**P*<0.01; \*\*\**P*<0.001). **(B-D)** Representative flow cytometric plots from each condition (IC; isotype control) following stimulation for 18 hours. **(B)** For IL-6 and IL-27 **(C)** For IL-12. **(D)** CD4^+^CRTAM^+^ T-cells in activated naïve CD4^+^ T-cells from *wt*, *Il6ra*^-/-^ and *Il27ra*^-/-^ mice. IL-6 or IL-27 was added as shown.

### Suppression of CRTAM expression on CD4*^+^* T-cells requires STAT3–

IL-6 and IL-27 signal *via* receptor systems sharing the common β-receptor subunit gp130. However, IL-27 promotes a stronger STAT1 signal, often inhibiting the action of other lymphokines^12,20^. To identify the signaling mechanisms responsible for CRTAM control by IL-6 and IL-27, we studied CRTAM expression on naïve CD4^+^ T-cells from *gp130*^Y757F:Y757F^ mice (**Fig-5**). These animals possess a single tyrosine-to-phenylalanine substitution in the cytoplasmic domain of gp130 that causes a prolonged hyperactivation of STAT1 and STAT3 following ligand activation^7,9,39,40^. Both IL-6 and IL-27 inhibited the expression of CRTAM on activated naïve CD4^+^ T-cells from *gp130*^Y757F:Y757F^ mice (**Fig-5A** & **B**). To identify the regulatory factors downstream of gp130, we analyzed CRTAM on naïve CD4^+^ T-cells from *gp130*^Y757F:Y757F^:*Stat3^+/-^* mice with genetically reduced gp130-dependent STAT3 activities^7,9,39,40^. Compared to the inhibition of CRTAM by IL-6 and IL-27 in activated naïve CD4^+^ T-cells from *gp130*^Y757F:Y757F^ mice, there was a marked reduction in the inhibitory profile of both cytokines in activated naïve CD4^+^ T-cells from *gp130*^Y757F:Y757F^:*Stat3^+/-^* mice (**Fig-5C**). These data point towards STAT3 as a repressor of CRTAM^+^CD4^+^ T-cells.

**Figure-5.**
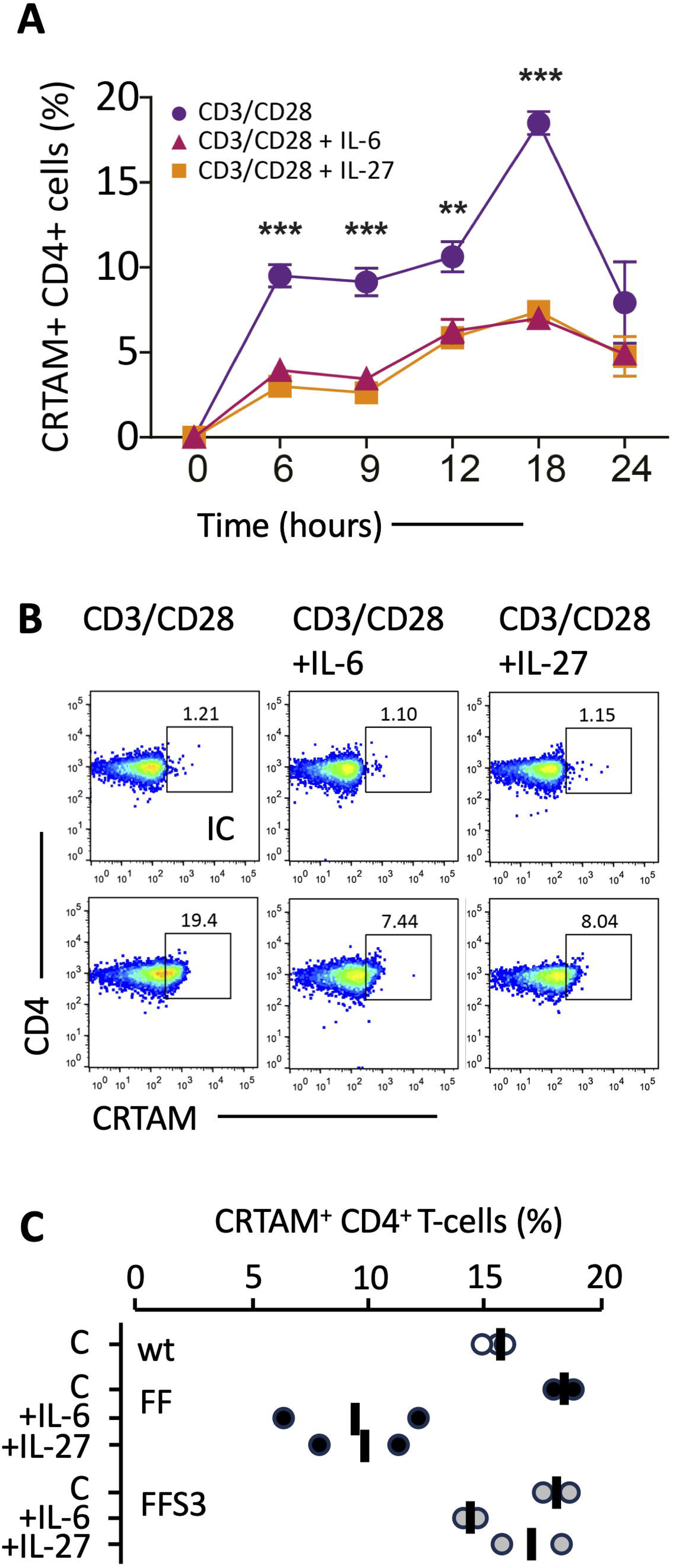
Jak-STAT cytokine signaling via gp130 controls CD4*^+^*CRTAM*^+^* T-cells. **(A)** Splenic naïve CD4^+^ T-cells from *gp130*^Y757F:Y757F^ mice were cultured with co-stimulatory anti-CD3 and anti-CD28 antibodies. Data shows CRTAM expression following treatment with 20 ng/ml IL-6 and 10 ng/ml IL-27. **(B)** Representative flow cytometry is shown for each experimental condition. **(C)** The proportion of CRTAM^+^CD4^+^ T-cells in activated splenic naïve CD4^+^ T-cells from *wt*, *gp130*^Y757F:Y757F^, and *gp130*^Y757F:Y757F^:*Stat3^+/-^* mice. Data show changes in CRTAM^+^CD4^+^ T-cells as a response to treatment with IL-6 or IL-27. Data was acquired from CD4 T-cells at 18 hours post-stimulation (mean ± SEM from 2 experiments; \**P*<0.05; \*\**P*<0.01; \*\*\**P*<0.001).

### ZEB1 regulates CRTAM expression by CD4*^+^*T-cells–

Computational methods (JASPAR) were used to identify putative DNA binding sites for STAT transcription factors in the *Crtam* promoter. Bioinformatic predictions failed to identify a tandem repeat element or a gamma-activated sequence (GAS) motif within the proximal promoter region for *Crtam* (**Fig-6A**). Instead, 46 paired CACCT(G) E-box-like sequences common to ZEB1 transcription factor binding were identified (**Fig-6B**). To consider the link between ZEB1 and CRTAM, we first examined their cellular expression in synovial tissues from mice with AIA and synovial biopsies from RA patients. This analysis identified the co-expression of ZEB1 and CRTAM in synovial infiltrating immune cells, including human CD4^+^ T-cells expressing CRTAM (**Fig-6C**). Immunofluorescent staining of synovial tissues from mice with AIA confirmed CRTAM co-localization with ZEB1 in synovial infiltrating cells (**Fig-6D**). Based on these initial observations, we asked if the STAT3 control of CRTAM required ZEB1. JASPAR identified 13 enhancer sites in the *Zeb1* promoter with homology with consensus sequences for STAT transcription factors (**Fig-6B**). We, therefore, used ChIP-qPCR to screen activated naïve CD4^+^ T-cells for STAT transcription factor binding to the *Zeb1* promoter (**Fig-7A**). Sequences containing GAS motifs from the promoters of *Irf1* and *Socs3* were used as controls for STAT binding (**Fig-7A**)^10^. ChIP-qPCR revealed STAT3 binding of the *Zeb1* promoter following IL-6 or IL-27 stimulation (**Fig-7A**). No enrichment of STAT1 was observed (**Fig-7A**). Despite efforts to determine the binding of ZEB1 to the *Crtam* promoter, we failed to identify an anti-ZEB1 antibody suitable for ChIP. We, therefore, sought to identify the relationship between *Zeb1* and *Crtam* in synovial tissues from mice with AIA (**Fig-7B**). Quantitative PCR showed that increases in synovial *Zeb1* coincided with decreases in *Crtam* expression. This inverse relationship was also observed in transcriptomic data from human synovial biopsies. Although synovial *ZEB1* expression was comparable in biopsies displaying fibroblast-driven, myeloid-driven, and lymphoid-driven synovitis, *ZEB1* levels correlated with increased *STAT3* in RA (**Fig-7C** & **D**). However, consistent with our mouse findings, we identified an inverse relationship between *ZEB1* and *CRTAM* expression in transcriptomic data from human synovial biopsies (**Fig-7E**). To test the role of ZEB1 in regulating CRTAM, we attempted to purify splenic naïve CD4^+^ T-cells from *Zeb1*^-/-^ mice^41^. Spleens from these mice showed gross morphological and cellular alterations in splenic architecture that hampered recovery of naïve CD4^+^ T-cells. Small numbers of splenic CD4^+^ T-cells were isolated from *Zeb1*^-/-^ mice and they showed no phenotypic markers (e.g., CD62L and IL-6R) of naïve CD4^+^ T-cells, which affected our ability to test the properties of Zeb1 in controlling CRTAM (**Supplemental Figure-2**). Nevertheless, our results show that CRTAM^+^ CD4^+^ T-cells are present during synovitis, with their involvement predominantly associated with lymphoid-rich synovitis.

**Figure-6.**
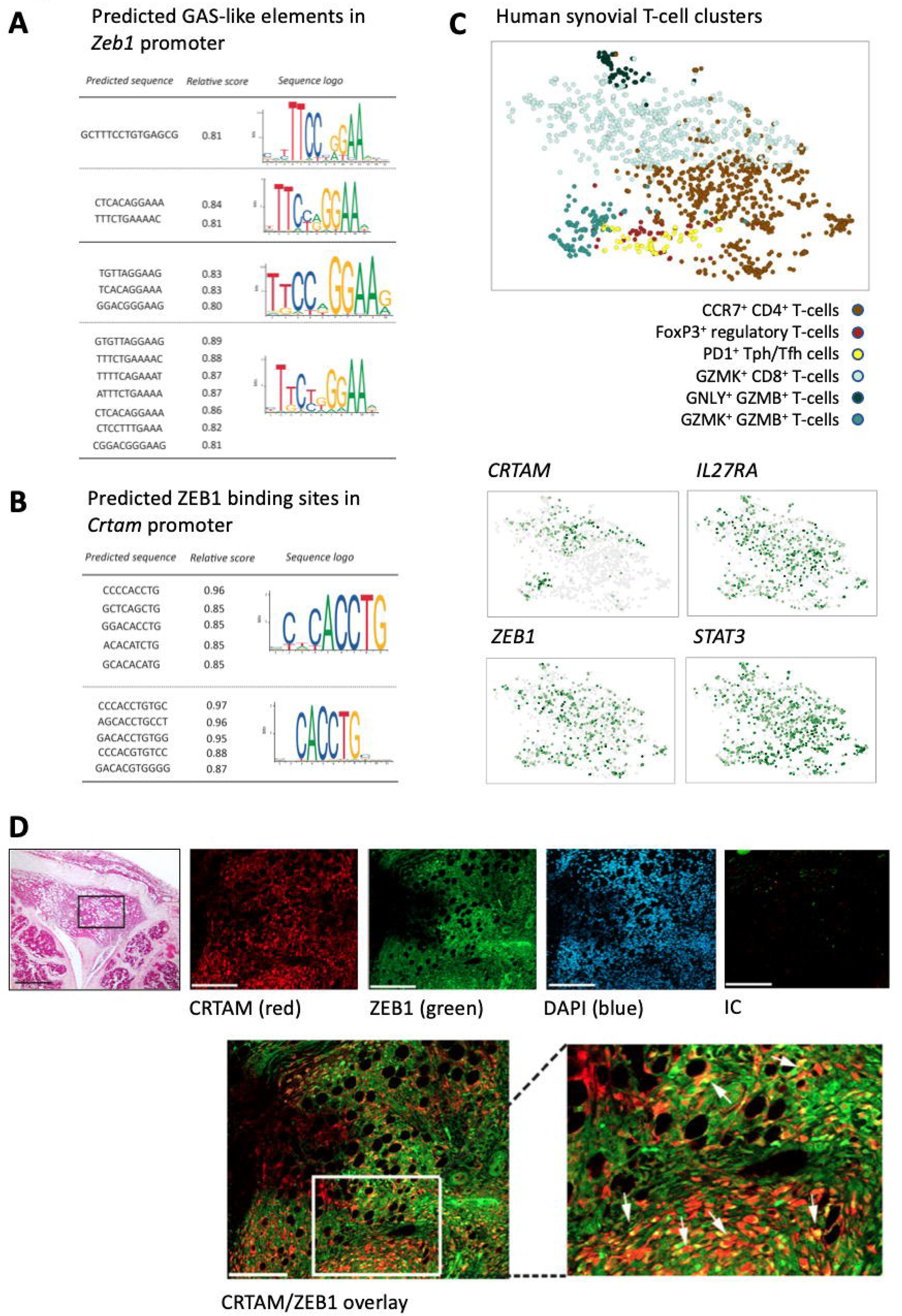
Co-expression of ZEB1 and STAT3 in active synovitis. **(A)** *In silico* analysis (JASPAR [http://jaspar.genereg.net/]) of putative transcription factor binding sites for STAT1 and STAT3 in the promoter of *Zeb1*. Sequence predictions show the sensitivity and specificity of each motif to consensus sequence annotation (upper panel). **(B)** Similar predictions for ZEB1 binding sites in the *Crtam* promoter (lower panel). **(C)** UMAP clustering of scRNA-seq data for human synovial T-cells from the *Accelerating Medicines Partnership* and mapped to T-cell subsets (upper panel)^43^. The localization of *CRTAM*, *IL27RA*, *ZEB1*, and *STAT3* is shown. **(D)** Representative image of H&E staining of mouse knee joints at day 10 post-AIA induction (scale bar, 500 µm). The boxed area locates the area visualized by immunofluorescence. Confocal fluorescent images are shown for CRTAM (red), ZEB1 (green), and DAPI nuclear counterstaining (blue) (scale bar, 100 µm). Isotype control staining is shown (IC). Synovial co-localization of CRTAM and ZEB1 staining is shown in the larger micrographs (yellow) and identified by the arrowheads.

**Figure-7.**
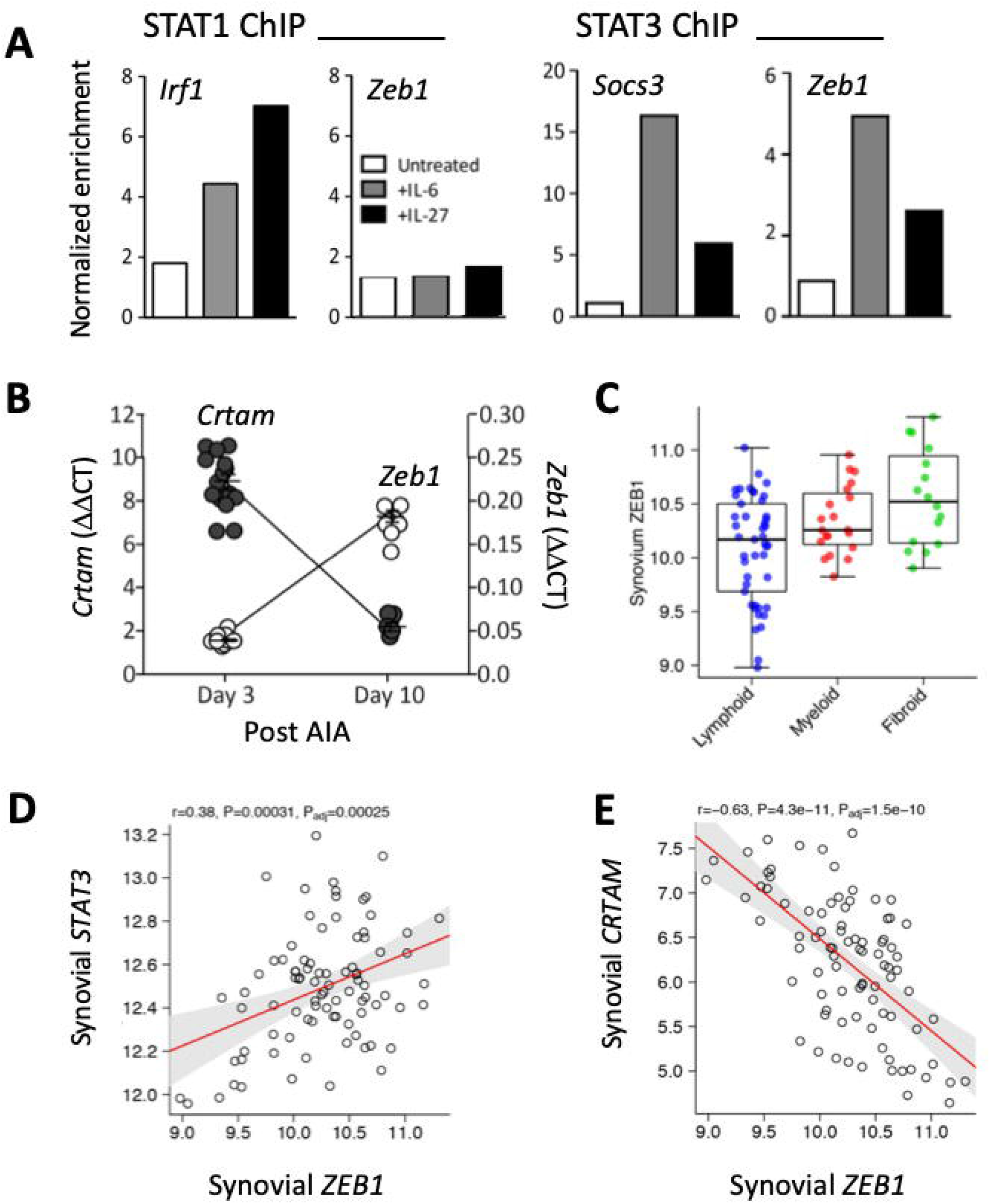
STAT3 control of CRTAM requires ZEB1. **(A)** ChIP-qPCR analysis of STAT1 and STAT3 binding to the *Zeb1* promoter using genomic DNA isolated from activated naïve CD4^+^ T-cells following IL-6 or IL-27 stimulation. STAT transcription factor binding to promoter sequences from *Irf1* (for STAT1) and *Socs3* (for STAT3) were used as controls. A single analysis was performed on genomic DNA pooled from naïve CD4^+^ T-cells extracted from 12 mice. **(B)** Quantitative PCR of *Zeb1* in synovial tissues from wt, *Il6ra*^-/-^ and *Il27ra*^-/-^mice with AIA (day-3 and day-10 of disease) (mean ± SEM; n=3-8 mice per group from three independent experiments; \**P*<0.05; \*\**P*<0.01; \*\*\**P*<0.001). **(C)** Analysis of RNA-seq data from synovial biopsies collected from the PEAC study. *ZEB1* expression in biopsies of known pathology is shown. **(D)** Pearson correlations of synovial *ZEB1* against *STAT3*. **(E)** Pearson correlations of synovial *ZEB1* against *CRTAM*.

## DISCUSSION–

Cellular interactions between infiltrating T-cells, the stromal compartment, and antigen-presenting cells support the maintenance of lymphocyte effector functions in inflamed tissues. Here, we report the identification of CRTAM^+^CD4^+^ T-cells in synovitis and describe a regulatory role for IL-6 and IL-27 in controlling CRTAM expression. CRTAM shares close homology with scaffolding proteins (e.g., DLG1, SCRIB, LGL) that facilitate the assembly of complexes required for lymphocyte adhesion, polarity, and proliferation. Our analysis showed CRTAM expression confined to a subset of synovial infiltrating lymphocytes. We identified a higher proportion of synovial CRTAM^+^CD4^+^ T-cells in exudates from *Il27ra*^-/-^ mice, where AIA promotes ELS formation^8^. Thus, CRTAM^+^CD4^+^ T-cells accumulate during active synovitis, with CRTAM potentially affecting cellular interactions with CD21 follicular dendritic cells or stromal fibroblasts. In support, we provide two pieces of evidence. First, CADM1, the natural ligand for CRTAM, was expressed by sub-lining fibroblasts involved in synovial immune regulation and preservation of cartilage integrity^43,45^. Second, synovial *CRTAM* correlates with effector cytokines (e.g., *IFNG*, *IL17A*, *IL21*, *IL22*) linked with lymphoid-driven synovitis. These same cytokine signatures resemble the effector properties of CRTAM^+^CD4^+^ T-cells involved in maintaining mucosal immunity^30,33,34^.

In contrast to the broad synovial expression of DLG1, SCRIB, and LGL on macrophages, fibroblasts, B-cells, and T-cells, CRTAM identifies with the effector phenotype of activated CD4^+^ T-cells in synovitis and correlates with the synovial expression of *GZMB*, *EOMES*, *IFNG*, and *CXCR3*. These markers define murine CD4^+^ T-cells displaying cytotoxic properties consistent with those of activated NK cells, NKT cells, and some T-cell subsets^23,25–28,30^. Considering the regulation of CRTAM on CD4 T-cells, we identified a potential involvement for lymphokines that use the gp130 cytokine receptor cassette. For example, IL-6 and IL-27 impact the activity or expression of granzymes, T-bet, IFNγ, perforin, and eomesodermin^46–48^. We now show that IL-27 is a negative regulator of CRTAM expression on activated naïve CD4^+^ T-cells, with *Il27ra*-deficient mice showing enhanced numbers of synovial infiltrating CRTAM^+^CD4^+^ T-cells following arthritis induction. This response is consistent with the immunomodulatory action of IL-27, which inhibits IL-2 production but induces IL-10 and immune checkpoints^12^. A similar blockade of CRTAM occurred with IL-6, and our findings describe a regulatory mechanism involving the STAT3 control of ZEB1, a member of the zinc-finger-homeodomain transcription factor family. Prior studies of ZEB1 bioactivity in T-cells identify that ZEB1 represses IL-2 and CRTAM expression and drives the induction of immune checkpoints^49–51^. These properties resemble those of IL-27, suggesting cooperation between gp130-activating cytokines and ZEB1 signaling. While this connection remains uncorroborated in T-cells, studies describe a link between these pathways in epithelial-to-mesenchymal transition and tumor metastasis in pancreatic cancer^52,53^.

Cytokine signaling through the Jak-STAT pathway regulates the recruitment of synovial infiltrating leukocytes^9,18,19^. Studies of experimental arthritis in gp130 knock-in mice epitomize this response, with the hyper-activation of STAT3 sustaining immune cell infiltration and promoting the organization of lymphoid aggregates in the inflamed synovium^7,9,21^. Our results show that increases in synovial CRTAM^+^CD4^+^ T-cells coincide with the incidence of synovial ectopic lymphoid-like structures. While the function of these cells requires further investigation, evidence of CRTAM involvement in gut mucosal immunity identifies links to the regulation of T-cell effector functions, Jak-STAT cytokine signaling in maintaining epithelial homeostasis, and Th17 responses to enteric pathogens^54,55^. Consistent with these findings, synovial tissues with evidence of ectopic lymphoid-like structures display gene signatures associated with IL-17, IL-21, or Th17 cells^8–11^. Thus, CRTAM signaling may control activities affecting organization, maintenance, or the function of synovial ectopic lymphoid-like structures. Additional research is required to understand the homing of CRTAM^+^CD4^+^ T-cells to CADM1-expressing stromal cells. Investigation of immune homeostasis in the gut suggests that CRTAM shapes the local inflammatory milieu to affect gut microbiota composition and response to intestinal pathogens^54^. This feature of CRTAM biology raises the possibility that CADM1 binding may promote cell adhesion and the generation of focal cytokine networks instructing lymphoid aggregation and organization into ectopic lymphoid-like structures.

## ACKNOWLEDGEMENTS–

The research was supported by grants from Versus Arthritis (Reference 20770, 19796, 20305), UKRI MRC (Reference MR/X00077X/1), Wellcome Trust (Reference 107964/Z/15/Z), and the National Health and Medical Research Council (NHMRC) of Australia. BJJ holds a Senior Research Fellowship from the NHMRC. Analysis of synovial tissues was conducted with support from the Medical Research Council (Reference 36661) and the Versus Arthritis-funded Experimental Arthritis Treatment Centres in QMUL and Cardiff (Reference 20022, 20016).

## Supplementary Figure Legends

**Supplemental Figure-1. The cytokine control of activated naïve CD4*^+^* T-cells.**

**(A)** Splenic naïve CD4^+^ T-cells from *wt* mice were cultured with co-stimulatory anti-CD3 and anti-CD28 antibodies. Temporal changes in CRTAM^+^ CD4^+^ T-cells are shown following treatment with 10 ng/ml IL-12. Representative flow cytometric plots are shown following stimulation for 18 hours. **(B)** Data analysis of publicly available transcriptomic datasets (E-MTAB-7682) from splenic naïve (nv) and activated naïve CD4^+^ T-cells (nv.act) (see reference 10 for experimental details). Expression intensity from the Affymetrix gene array is shown for CD4^+^ T-cells treated with 20 ng/ml for both conditions (nv.IL-6 and nv.act.IL-6). The table shows an analysis of a select set of genes displaying both a relative signal intensity of >150 and >1.5-fold suppression in expression following IL-6 treatment (P<0.05). The regulation of *Socs1* is shown as a positive control for IL-6 signaling.

**Supplemental Figure-2. Phenotypic properties of splenic lymphocytes from *Zeb1^-/-^* mice.**

**(A)** Photographic images of spleens isolated from *wt* (right) and *Zeb1*^-/-^ (left) mice. **(B)** Splenic CD4 T-cells were pooled from 3 mouse spleens. Representative FACS analysis of CD4^+^ T-cells from *wt* and *Zeb1*^-/-^ mice, with the histograms showing staining for CD44, CD62L and IL-6R. **(C)** Naïve CD4^+^CD25^-^CD44^lo^CD62L^hi^ T-cell numbers from *wt* and *Zeb1*^-/-^ (KO) mice. Values show cell numbers from two independent experiments using splenic CD4 T-cells pooled from 3 mouse spleens per genotype.

