## Supplementary figures and images for "IL-6 and IL-27 negatively regulate CRTAM-expressing CD4^+^ T-cells associated with lymphoid-driven synovitis"

### Supplementary Fig.1

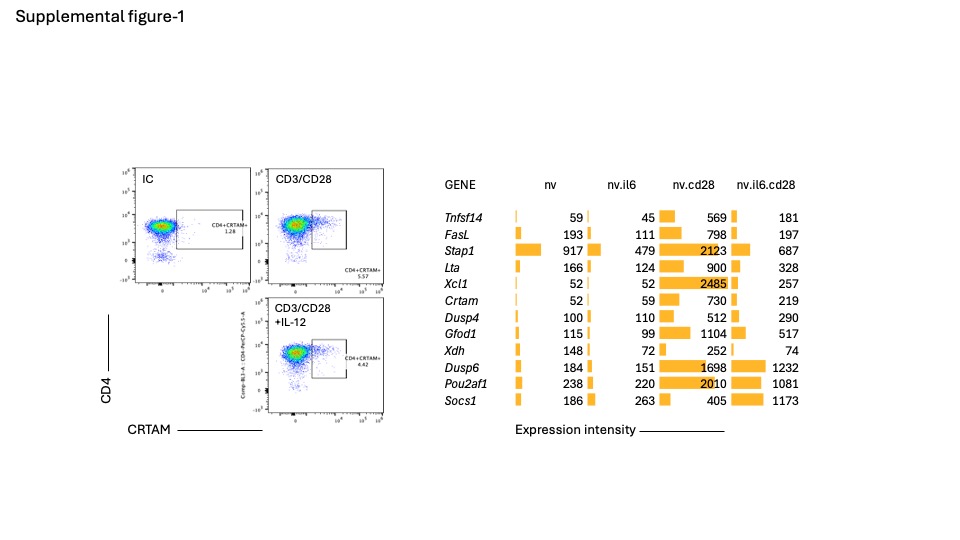

### Supplementary Fig.2

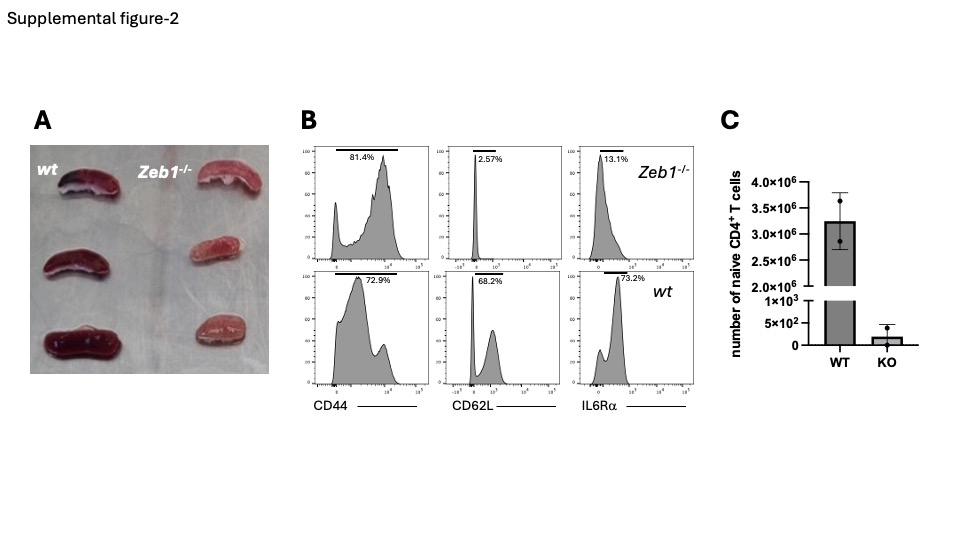
